# Potential clinical benefits of CBD-rich Cannabis extracts over purified CBD in treatment-resistant epilepsy: observational data meta-analysis

**DOI:** 10.1101/212662

**Authors:** Fabricio A. Pamplona, Ana Carolina Coan

## Abstract

**Potential clinical benefits of CBD-rich Cannabis extracts over purified CBD in treatment-resistant epilepsy: observational data meta-analysis**

Different therapies involving cannabinoid compounds have become popular on the past few years, particularly the use of canabidiol (CBD) based products for the treatment of child refractory epilepsy. In this segment we highlight genetic disorders such as the Dravet Syndrome, which has received a lot of attention in Brazil. Our country has been giving visibility for this issue after the public discussion regarding the patient Anne Fischer, who benefited from a treatment with hemp based products, imported from the United States of America as a nutritional supplement, and still unregistered in Brazil. To this moment there is no cannabinoid based product registered for clinical indication for epilepsy, so patients have at their disposal products considered nutritional supplements, in general produced from a type of Cannabis known as “hemp”, which are commercialized in Brazil through medical prescription. Despite several anecdotal evidences from patients and family members, until now there is no consensus on medical literature over the efficacy and safety of these products. Some observational studies are available on scientific literature, but there is still a scarcity of clinical studies conducted under the logic, rigor and organization necessary for clinical trial dedicated to the register of a pharmaceutical product. The objective of this paper is to describe the analysis of several observational clinical studies available on the literature regarding the treatment of child refractory epilepsy with cannabinoid based products. Beyond attempting to establish the safety and efficacy of such products, when possible, the present analysis also intended to investigate if there is enough evidence between the different aspects of safety and efficacy between CBD enriched extracts compared to purified CBD products. Results: a systematic search for papers in the “PubMed” search system with the words “Dravet”, "Lennox-Gastaut" and "epilepsy" combined with the terms "Cannabis", "cannabinoid" and "child" yielded 30 papers. From those, 24 were not considered for the systematic review, for not having valid content (13), for being opinion only papers (6), showed not clinical data (4) and reported different subjects (1), resulting in 6 valid papers published between 2013 and 2016. One additional study was *in press* at the moment of the search was added manually (1) resulting in 7 valid references for analysis, with an average impact factor of 5,9 (2,3 to 21,8). Public data from partial reports of controlled randomized studies conducted in order to register a medication (2) were also considered, when appropriated and were mentioned in the text. The categorical data were analyzed by the Fischer test. Overall, the papers analyzed report observational clinical data of 442 patients, treated with CBD rich extracts or purified CBD, with he average daily dose between 1 and 50 mg/kg, with treatment length from 3 to 12 months (average of 6,2 months). A considerable amount of 66% (292/442) of the patients reported improvement in the frequency of convulsive crisis. There were more reports of improvement from patients treated with purified CBD (242/285) than patients treated with purified CBD (68/157), with statistical significance (p<0,0001). Nevertheless, when the standard clinical threshold of a “50% reduction or more in the frequency of convulsive crisis” was applied, only 40% of the individuals are considered respondent, and there were no difference (p=0,57) between the treatments with extract (64/168) and purified CBD (65/157). However, even that both treatments have similar efficacy, the patients treated with CBD enriched extracts reported a lower average dose than purified CBD patients. The average CBD equivalent dose on the extracts was 7,1 mg/kg/day, while the purified CBD was 22,9 mg/kg/day, suggesting that CBD is about 3x more potent in the extract than in its purified form. Looking only at the data relative to genetic originated disorders, there is evidence of a superior efficacy on Dravet Syndrome patients (37/72, p=0,01), but not for Lennox-Gastaut Syndrome (78/188, p=0,18), compared to the number of refractory epilepsy respondents in general (107/305). There is also an advantage of the CBD enriched extracts related to the occurrence of side effects. The report of mild side effects (109/285 vs. 291/346, p<0,0001) and severe (23/285 vs. 77/346, p<0,0001) are more frequent in products containing purified CBD than on CBD enriched extracts. Important to mention that these are the numbers of total reports of side effects, it is not possible to infer which fraction of these numbers are related to the treatment. In conclusion, this meta analysis suggests that treatments using CBD enriched extracts are more potent and have a better profile of adverse effects (but not more efficacy) than products containing purified CBD, at least in this population of patients with refractory epilepsy. The lack of standardization between extracts containing Cannabis does not allow us to infer directly which characteristics of the product that confer this therapeutic advantage, but it is likely related to other compounds present in the formulation that act sinergistically with CBD. Controlled studies with standardized Cannabis based extracts are necessary to confirm these observations.

\**Presented as an abstract and lecture to the 2017 CannMed event, at the Harvard Medical School, Boston, USA, in may 2017.*

## 1 Introduction

Different therapies including Cannabinoid compounds became popular in the past 2 years in Brazil, particularly the use of canabidiol (CBD) based products for the treatment of child refractory epilepsy. In this segment, we highlight genetic disorders such as the Dravet Syndrome, that has been receiving a lot of attention in Brazil. Our country has been giving more visibility for this subject after the public discussion about the case of the patient Anne Fischer, who benefited from a hemp based product treatment, imported from the US as a nutritional supplement, and still unregistered in Brazil. From this symbolic case, it was established a mechanism of facilitated importation of this type of product, which has already benefited around three thousand patients until may 2017 (Figure 1). The growth projection of the need for prescriptions for imported products is exponential, with a growth rate of 85% per year, passing the landmark of 15 thousand special authorizations to import by the beginning of 2020. That is considering only this mechanism, which even being facilitated by the regulatory agency, still continues to be very bureaucratic and allows the access to products considered expensive and, in many cases, with low credibility, given the suspicion of low quality and the absence of efficacy and safety confirmation. So, it is important that we rely on data available until this moment in an attempt to provide more legitimacy to the treatments that have been realized, in benefit of greater safety to patients, their families and prescribing physicians.

**Figure 1.**
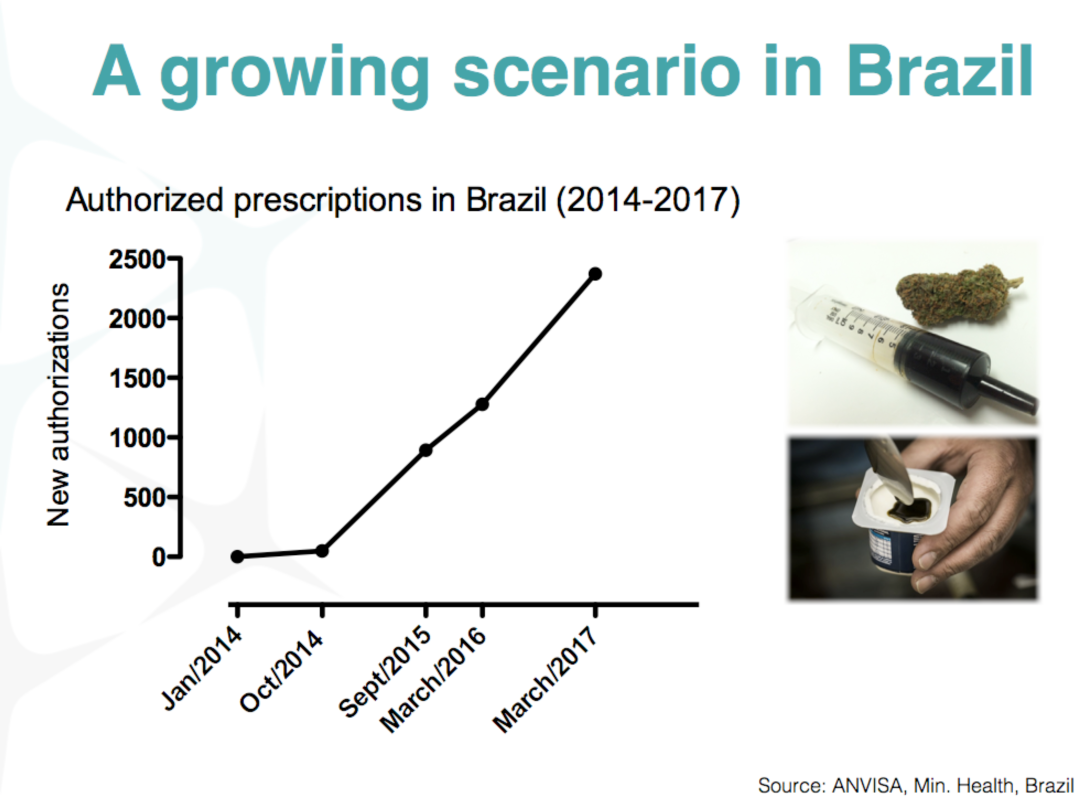
Data of patients with special authorization to import Cannabis based products in Brazil, between january 2014 and march 2017.

To this moment there is no cannabinoid based product registered for the treatment of epilepsy in the world, in a way that these patients have at their disposal products considered nutritional supplements, in general prepared from a type of Cannabis known as “Hemp”, and commercialized in Brazil under a physician’s prescription. These products are not considered narcotics at the original place of their production, and are being distributed to Brazil under an exception regime. The real need of the patients will only be met with the availability of products supported by robust clinical data that supports its efficacy and safety, but until this moment, such product is not available anywhere in the world. That is the specific void that Entourage Phytolab’s first product intends to fill, with clinical indication of child refractory epilepsy.

Despite several anecdotal evidences of patients and family members, broadly publicized in the country though several communication outlets, until now, there is no consensus on the medical literature about the efficacy and safety of these products. Some observational studies are available on scientific literature, but there is a scarcity of clinical data acquired within the logic, rigor and organization necessary to the conduction of clinical studies destined to the registration of a pharmaceutical product.

The objective of the present paper is to describe an analysis of clinical observational studies available on the literature for treatment of child refractory epilepsy with the use of cannabinoid based products, in a way that it subsidizes the definition of safety and efficacy of this therapeutic approach. Beyond attempting to establish the safety and efficacy of these products, when possible, the present analysis also intended to investigate if there is evidence of differences in the aspects of safety and efficacy between CBD enriched extracts compared to purified CBD products.

## 2 Meta analysis search strategy

### 2.1 Identification, inclusion and exclusion

A systematic search was made on the databases of MEDLINE/PubMed (http://www.ncbi.nlm.nih.gov/pubmed) and Google Scholar (http://scholar.google.com) intending to identify original papers with observational or experimental data about the use of Cannabis and its compounds on the treatment of refractory epilepsy in humans. The search was limited to papers published in English, with results obtained from human beings. The titles, abstracts and full texts of all search results had their eligibility analyzed, considering inclusion and exclusion criteria. Inclusion criteria: studies containing clinical data in humans, that could infer the efficacy and safety profile of the products containing cannabinoids for epilepsy. Exclusion criteria: Review and opinion papers, case studies, studies with no measurable data, studies wit no accessible numerical data. Were considered eligible papers describing studies in observational, prospective and retrospective format, other than controlled studies, regardless of the kind and duration of the treatment.

### 2.2 Definition and treatment of clinical data

Classically utilized objective clinical outcomes were defined on the research fin epilepsy to group the studies on a meta analysis format. Were considered the clinical outcomes of “improvement reports” and “reduction of the number of crisis”. This second, considered in two thresholds, the reduction of the frequency of crisis above 50% (considering individuals “respondent” to the treatment) and the reduction of frequency of crisis above 70%. As clinical safety outcomes, were considered all reported outcomes in the papers as adverse effects. The objective data considered was the frequency of the report of adverse events, distinguished as “mild” and “severe”, such as written in every paper.

### 2.3 Definition of variables and statistical analysis

The data on every paper was grouped under the format of categorical variable (the number of patients in regard of the group). The cases in which the paper describes data in a format of continuous variable (percentage, or individual frequency reduction), the data was inferred/estimated and transformed in categorical variable for further analysis. The transformed data was analyzed statistically by the Fischer test for categorical variables.

## 3 Clinical studies considered in the meta analysis

The systematic search took place in 02/09/17 using the key-words “epilepsy”, “Dravet”, “Lennox-Gastaut” combined with "Cannabis", "cannabinoid" and "child" resulting in 30 papers. From these, 24 were not considered for the systematic review, because they had no valid content (13), they were opinion papers (6), they reported non-clinical data (4) or reported other pathologies unrelated (1), resulting in six valid papers, according to the daygram demonstrated in the Figure 2 below. One additional study was in press at the moment of the search was added manually (1) resulting in 7 valid references for analysis, with an average impact factor of 5,9 (2,3 to 21,8). Public data from partial reports of controlled randomized studies conducted in order to register a medication (2) were also considered, when appropriated and were mentioned in the text.

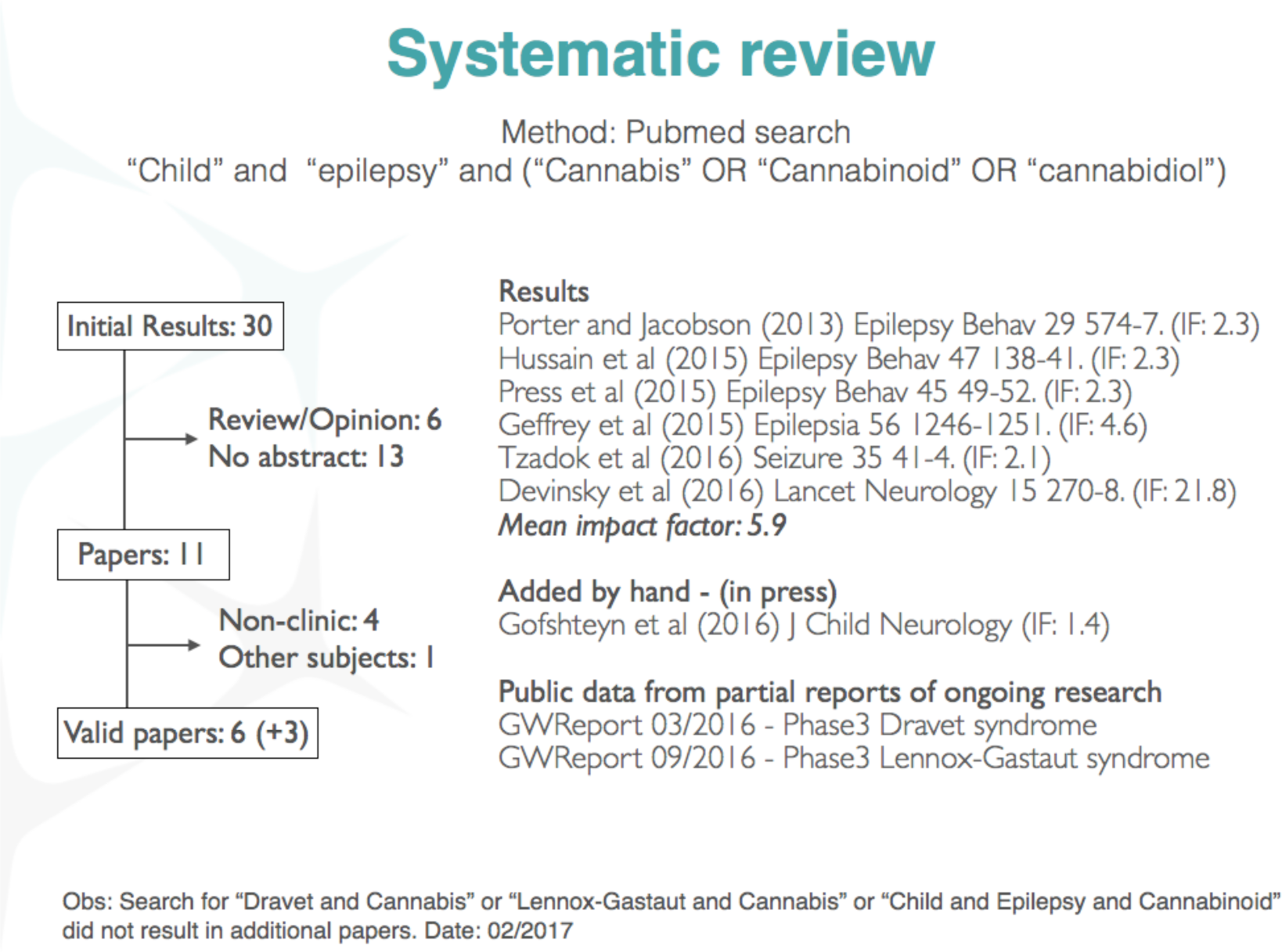

The studies included are fairly recent, published between 2013 and 2016, demonstrating how vivid this subject is in the world literature. Overall, the papers analyzed report observational clinical data from 442 patients, treated with CBD enriched extracts or purified CBD, with average daily doses between 1 and 50 mg/kg, with duration of treatment from 3 to 12 months (average of 6,2 months), as shown in the Table below.

From the studies selected to analysis, 4 show a retrospective design (with a total of 285 patients) and 3 show a prospective design (with a total of 157 patients). The quality of the evidences shown varies, being that 2 studies used research based on online questionnaires with family members and care takes (with a total of 136 patients) and 5 studies used evidences of improved quality, originated from medical history (with a total of 231 patients). As for the type of treatment, 3 studies report data from patients who used purified CBD (total of 157 patients) and 4 studies report data from patients who used herbal extracts rich in CBD, of different nature (total of 285 patients). It doesn’t seem to have evidences of an obvious bias that compromises the interpretation of data, considering the format of data gathering of each study. The only reservation is that all studies which used purified CBD had a prospective data gathering, while all studies which used CBD enriched extracts had a retrospective data gathering. All studies used a heterogeneous population of epilepsy patients, and the segmentation in specific types of syndromes was done afterwards. All studies were conducted by medical centers experienced in conducting this type of study, at universities or internationally reputed research centers. Curiously, 6 out of the 7 studies were conducted or leaded (when it was a multicentric study) by universities or research centers in the United States. One study was conducted in Isreal.

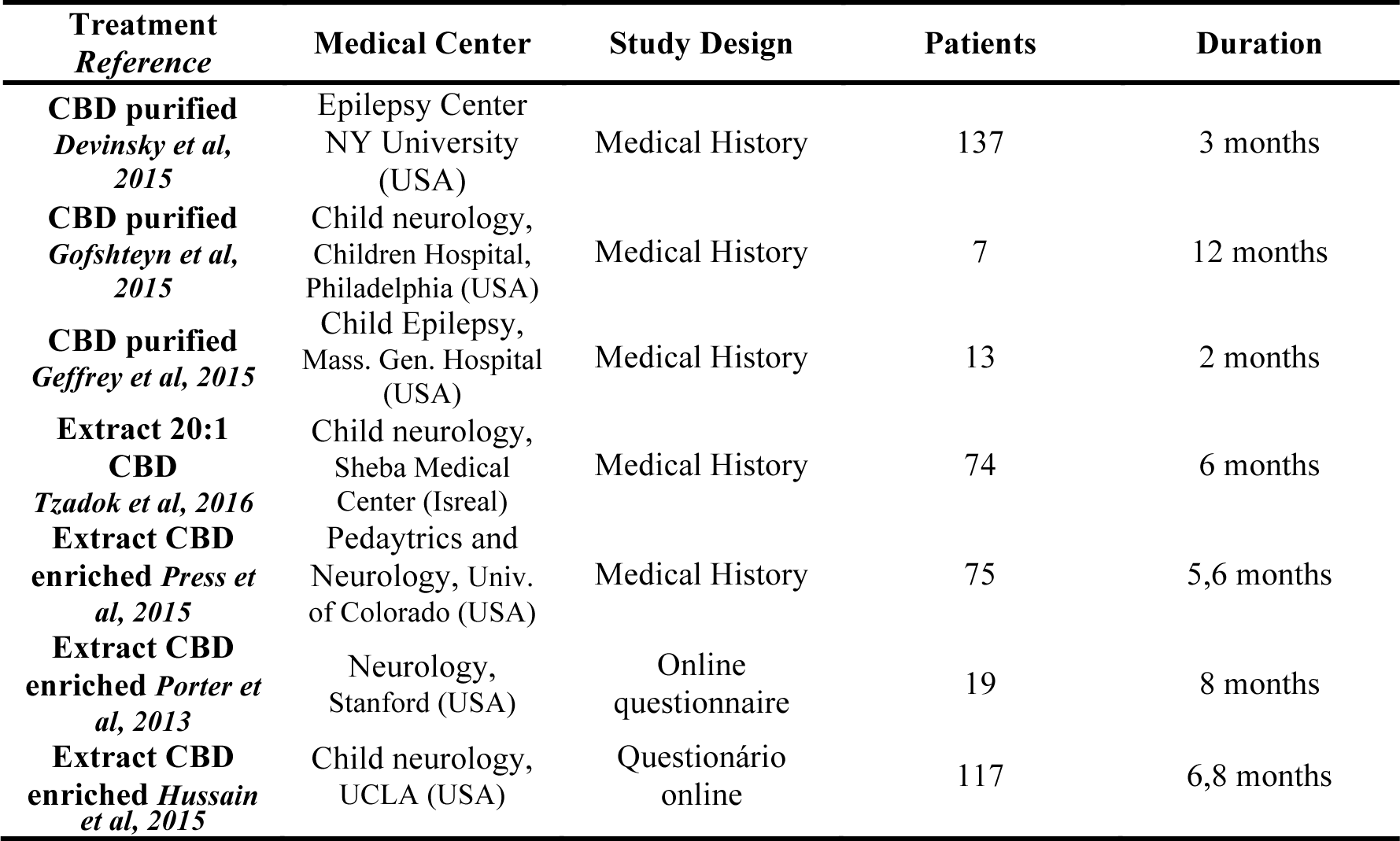
Table XX.

The studied population consisted in children and adolescents, between 1 and 18 years old with treatment resistant epilepsy (refractory epilepsy), who tried between 4 and 12 other medications for 3 years before trying canabidiol based products (CBD). That is, it is a very refractory population, of difficult treatment. Most of the patients is carrier of an epileptic encephalopathy, highlighting a percentage of patients with Dravet and Lennox-Gastaut syndromes, adding up to 26% of the total patients, which were evaluated separately on a future analysis.

## 4 Discussion of the results

The general results allow the affirmation that the treatment with CBD based products improve the symptoms associated to epilepsy, even for this refractory population. A considerable fraction of the patients reported an improvement in the frequency of convulsive crisis, 66% (292/442), with the studies varying between 37% and 89%. Five out of seven studies showed over 80% of the patients reporting improvement. There were more reports of improvement per treated patient with CBD enriched extracts (224/285, 78%) than those treated with purified CBD (68/157, 43%), with statistical significance (p<0,0001). However, when the clinical threshold of “50% or greater reduction in the frequency of convulsive crisis” was applied, “only” 40% of the individuals are considered respondent (studies varying from 33% to 74%), and there is no difference (p=0,57) between treatments with CBD enriched extracts (64/168, 38%) and purified CBD (65/157, 41%). Around one quarter of all treated patients (24%) are unresponsive for the endpoint “improvement greater than 70%” (the studies vary between 18% and 57% of respondents in this category), some studies report seizure free patients. The “seizure free” endpoint was not used for the analysis because it is not a parameter utilized by a significant amount of the studies. It is possible that the number of individual free of crisis to be around 15% to 20%, which is absolutely amazing for a population such as this, that has been testing without success several anti-epileptic medications, in mono therapy or combined. (Table XX).

Table XX. Efficacy of treatments in the reduction of convulsive crisis (heterogeneous population). Endpoints of crisis improvement, improvement greater than 50% (“respondent”) and greater than 70%, and average dose reported.

| **Treatment**<br>**Reference** | **Patients** | **Reported**<br>**improvement** | **>50%** | **>70%** | **Average daily**<br>**dose**<br>**(mg/kg/day)** |
| --- | --- | --- | --- | --- | --- |
| Total | 442 | 292/442 | 129/325 | 61/250 | (2mg/kg -<br>50mg/kg) |
| **Average** | **100%** | **66%** | **40%** | **24%** | **15mg/kg** |
| **CBD purified**<br>**Devinsky et al,**<br>**2015** | 137 | 37% | 37% | 22% | 22,9mg/kg |
| **CBD purified**<br>**Gofshteyn et al,**<br>**2015** | 7 | 86% | 71% | 57% | 22mg/kg |
| **CBD purified**<br>**Geffrey et al,**<br>**2015** | 13 | 85% | 70% | 46% | 24,6mg/kg |
| **Extract 20:1**<br>**CBD**<br>**Tzadok et al,**<br>**2016** | 74 | 89% | 34% | 18% | <10mg/kg |
| **Extract CBD**<br>**enriched Press**<br>**et al, 2015** | 75 | 57% | 33% | X | X |
| **Extract CBD**<br>**enriched**<br>**Porter et al,**<br>**2013** | 19 | 84% | 74% | 42% | 7mg/kg |
| **Extract CBD**<br>**enriched**<br>**Hussain et al,**<br>**2015** | 117 | 85% | X | X | 4,3mg/kg |

Curiously, even when both treatments had similar efficacy, when considered the classic parameter of “reduction greater than 50%”, the patients treated with CBD enriched extracts reported a lower average daily dose than purified CBD patients. The average daily dose for purified CBD was 22,9mg/kg/day, while the average daily dose of CBD equivalent reported for the treatment with CBD enriched extract was 7,1mg/kg/day. This data suggests CBD to be 3x more potent when administered in herbal form, probably suggesting that other minor compounds present in the extract may contribute for the therapeutic effect.

Looking only at the data relative to disorders with genetic origin (epileptic encephalopathy), considering individuals with an improvement greater than 50% (“respondents”), there is evidence of a superior efficacy on Dravet Syndrome patients (37/72, 51%, p=0,01), but not for Lennox-Gastaut Syndrome (78/188, 41%, p=0,18), compared to the number of refractory epilepsy respondents in general (107/305, 35%). This difference in response between such pathologies is confirmed by recent controlled randomized studies conducted by the company GW Pharma in patients with Dravet and Lennox-Gastaut Syndrome, where there is a greater difference between the average reduction of crisis compared to placebo on patients with Dravet Syndrome (39% vs 13% placebo) than the same parameter observed for the Lennox-Gastaut Syndrome population (37-42% vs 17% placebo) (GW Report 03/2016 e GW Report 09/2016).

Table XX - Comparison between the number of respondents with a reduction greater than 50% of the crisis between genetic encefalopathies.

Table XX - Comparison between the number of respondents with a reduction greater than 50% of the crisis between genetic encephalopathies.
| **Treatment**<br>**Reference** | **Patology** | **Respondents**<br>**(crisis**<br>**improvement >**<br>**50%)** | **%** |
| --- | --- | --- | --- |
| *Devinsky + Tzadok +*<br>*Press + Porter* | **Refractory Epilepsy,**<br>Combined studies | **107/305** | **35%** |
| Extract 20:1 CBD<br>*Tzadok et al, 2016* | **Epileptic encephalopathy** with or without<br>genetic confirmation* | **23/45** | **51%** |
| CBD pure<br>*Devinsky et al, 2015* | Dravet Syndrome | 16/32 | 50% |
| Extract c/ CBD<br>*Press et al, 2015* | Dravet Syndrome | 3/13 | 23% |
| Extract c/ CBD<br>*Porter et al, 2013* | Dravet Syndrome | 7/12 | 58% |
| Extract c/ CBD<br>*Hussain et al, 2015* | Dravet Syndrome | 11/15 | 73% |
| **Respondents Dravet** |  | **37/72** | **51%** |
| CBD pure (LG)<br>GW report 09/2016 | Lennox-Gastaut Syndrome <sup>\$</sup> | 59/149 | 40% |
| CBD pure<br>*Devinsky et al, 2015* | Lennox-Gastaut Syndrome | 11/30 | 37% |
| Extract c/ CBD<br>*Press et al, 2015* | Lennox-Gastaut Syndrome | 8/9 | 89% |
| **Respondents Lennox-Gastaut** |  | **78/188** | **41%** |

The specific population of each syndrome is too small to allow us to confidently infer regarding the differences between product type, but in general, we can consider that the CBD treatment has positive effects in such patients, regarding the reduction in frequency of the convulsive crisis.

Beyond the direct therapeutic effect of CBD in reducing epileptic crisis, reports about improvement in “secondary” aspects are very common, that give a great gain in quality of life for the patients and their family members. These secondary endpoints were reported by 285 patients. Perhaps we can infer that about 65% of the individuals who were treated obtained secondary improvements, but the truth is that this number might be considerably greater, since not all studies considered such endpoints during their development. The main “secondary” gains found were increased awareness (147/285, 52%), gains in quality of sleep (88/285, 31%), mood improvement (87/285, 30%), behavioral improvements (56/285, 20%), language use improvements (19/285, 7%) and improved motor skills (19/285, 7%) (Table XX).

Table xx - Secondary endpoints improvement reports, not directly associated to the reduction in epileptic crisis.

| **Treatment**<br>**Reference** | **n** | **Awareness** | **Sleep** | **Mood** | **Behavior** | **Language** | **Motor** |
| --- | --- | --- | --- | --- | --- | --- | --- |
| Extract 20:1 CBD<br>*Tzadok et al, 2016* | 74 | 34% | 11% | NR | 34% | 15% | 15% |
| Extract c/ CBD<br>*Press et al, 2015* | 75 | 33% | 7% | NR | 33% | 11% | 11% |
| Extract c/ CBD<br>*Porter et al, 2013* | 19 | 74% | 68% | 79% | 32% | NR | NR |
| Extract c/ CBD<br>*Hussain et al, 2015* | 117 | 71% | 53% | 63% | NR | NR | NR |
| Secondary endpoint<br>reported improvement | 285 | 147/285 | 88/285 | 87/285 | 56/285 | 19/285 | 19/285 |
| **Average** | **100%** | **52%** | **31%** | **30%** | **20%** | **7%** | **7%** |

There were also reports of other improvements, but we are considering only the ones which affected at least 5% of the studied population. Probably these effects occurred because of the reduction of the crisis, but in many cases they occurred before or even in the absence of significant reductions of convulsive crisis. There were no reports in secondary aspects in studies utilizing purified CBD (Geffrey et al, 2015; Gofshteyn et al, 2015; Devinsky et al, 2015), but it is not possible to conclude that no improvements on secondary endpoints occur with this type of treatment; it is more likely that the study wasn’t focused on this clinical aspect. In this way, analysis of adverse events considered only 285 individuals treated with CBD enriched extracts, according to Table XX. As demonstrated in one of the studies with 117 patients (Hussain et al., 2015), in a direct comparison of the same population, the conventional anti epileptics cause an improvement in these “secondary” parameters related to quality of life, but this effect occur in a smaller scale than in CBD based treatments. This is true, at least, for the effects in mood improvement, awareness, sleep quality and self-control. This data suggests that secondary positive events described for CBD are characteristic of the effects of this substance, and not only due to the reduction in the frequency of convulsive crisis (Table XX).

Table XX - Positive secondary adverse events of CBD and conventional anti epileptics. Reproduced from Hussain et al, 2015.

| Secondary Positive Effects |  |  |
| --- | --- | --- |
| Adverse Effects | Canabidiol | Anti-epileptics |
| Mood | 15/19 (79%) | 4/22 (18%) |
| Awareness | 14/19 (74%) | 6/22 (27%) |
| Sleep quality | 13/19 (68%) | 5/22 (23%) |
| Self-control | 6/19 (32%) | 3/22 (14%) |
| **Average** | **63.2%** | **20.5%** |

Although it is considered very safe in comparison to other anti epileptics, the treatment with CBD products is not free of adverse effects. The studies mentioned the occurrence of adverse events on a relatively large portion of the population studied (217/422, 51%), even though the great majority of events are considered “mild”. Severe events were reported by a much smaller portion of patients (64/422, 15%). In this case are being considered only patients of studies who mentioned the occurrence of adverse effects; if the study did not mentioned adverse events, we considered that it is not reasonable to assume that there were no adverse events and therefore we excluded the study of this analysis as a whole. Two studies containing only 20 patients were excluded according to this criteria (Geffrey et al, 2015; Gofshteyn et al, 2015). Important to mention that these are the total adverse effects numbers, it is not possible to infer which fraction of these are related to the treatment. The data is summed at Table XX.

Table XX - Reports of adverse events.

Table XX - Reports of adverse events.
| **Treatment**<br>**Reference** | **n** | **Mild/Moderate** | **Severe** |
| --- | --- | --- | --- |
| CBD pure<br>*Devinsky et al, 2015* | 137 | 79% | 30% (12%) |
| Extract 20:1 CBD<br>*Tzadok et al, 2016* | 74 | 46% | 18% |
| Extract c/ CBD<br>*Press et al, 2015* | 75 | 44% | 13% |
| Extract c/ CBD<br>*Porter et al, 2013* | 19 | 37% | 0% |
| Extract c/ CBD<br>*Hussain et al, 2015* | 117 | 30% | 0% |
| Total reports | 422 | 217/422 | 64/422 |
| **Average** | **100%** | **51%** | **15%** |

Curiously, there is also an advantage of CBD enriched extracts in relation to purified CBD regarding to the occurrence of adverse events. The reports of mild adverse effects (109/285 vs 291/346, p<0,0001) and severe (23/285 vs 77/346, p<0,0001) are more frequent in products containing CBD purified that in CBD enriched extracts. So we can make this comparison, since there were no reports of adverse events in the observational studies, were also included the adverse events reported in the controlled randomized clinical tests, according the report of the sponsor company (Table XX).

| **Treatment**<br>**Reference** | **n** | **Mild/Moderate** | **Severe** |
| --- | --- | --- | --- |
| CBD pure (SD) <sup>§</sup> | 120 | 50/60 | 10/60 |
| GW report 03/2016 | (2 branches) | 84% | 17% |
| CBD pure (LG) <sup>§</sup> | 255 | 133/149 | 26/149 |
| GW report 09/2016 | (3 branches) | 84-94% | 17% (5%) |

Table XX below describes the adverse events reported by the studies on a portion equal or greater than 5% of the individuals, considered “common”, and segmented in mild and severe. The most common events were the alteration of appetite, sleepiness, gastrointestinal disturbances/diarrhea, weight changes, fatigue and nausea. Uncommon or rare adverse events include thrombocytopenia, respiratory infections and alteration of the liver enzymes. There was a worsening of the convulsive state in some cases, but that cannot be attributed to the treatment. Uncommon or rare events reported occurred in combination with other anti-epileptic medication, particularly valproic acid and Clobazam, and might be related to drug interaction, and not to CBD toxicity.

Table XX - Adverse events reported in the studies.

Table XX - Adverse events reported in the studies.
| Reference | (Sample) | Freq. | AE mild common (>5%) | AE severe common (>5%) |
| --- | --- | --- | --- | --- |
| CBD pure<br>*Devinsky et al, 2015* | Medical records<br>prospec. (n=137) | 79%<br>(30%<br>graves) | Sleepiness (25%), Appetite<br>reduction (19%), Fatigue (13%),<br>Appetite increase (9%), Lethargy<br>(7%), Weight gain (7%), Weight<br>loss (6%) | Worsen convulsions (11%), Status<br>epilepticus (8%), Diarrhea (19%),<br>Hepatotoxicity (7%) |
| CBD pure (LG) <sup>§</sup><br>GW report 09/2016 | RCT vs placebo<br>(n=225) | 84-94% | Sleepiness, Appetite reduction,<br>Vomit, pyrexia e nasofaringite | Respiratory infection, Diarrhea e<br>epileptic status |
| CBD pure (SD) <sup>§</sup><br>GW report 03/2016 | RCT vs placebo<br>(n=120) | 84% | Sleepiness, Fatigue, Appetite<br>reduction, pyrexia, Vomit,<br>Lethargy | Respiratory infection, Diarrhea e<br>Convulsion |
| Extract 20:1 CBD<br>*Tzadok et al, 2016* | Medical records<br>retrosp. (n=74) | 46% | Sleepiness/Fatigue (22%)<br>Discomfort TGI (7%) | Worsen convulsions (18%) |
| Extract c/ CBD<br>*Press et al, 2015* | Medical records<br>retrosp. (n=75) | 44% | Sleepiness/Fatigue (12%)<br>Discomfort TGI (11%)<br>Irritability (5%) | Worsen convulsions (13%) |
| Extract c/ CBD<br>*Porter et al, 2013* | Online questionnaire<br>(n=19) | 37% | Dizziness (37%),<br>Fatigue (16%), Redução Appetite<br>(5%) | No reports of severe AE |
| Extract c/ CBD<br>*Hussain et al, 2015* | Online questionnaire<br>(n=117) | 30% | Appetite gain (30%), Weight gain<br>(29%), Dizziness (13%),<br>Irritability (9%),<br>Appetite reduction (6%),<br>Nausea (6%) | No reports of severe AE |

In order to verify this hypothesis, it is convenient to observe the data mentioned on one of the studies with 117 patients (Hussain et al, 2015), where the occurrence of adverse effects was analyzed before and after the treatment with CBD. Before the treatment with CBD, 36,9% of the patients reported in average 5 adverse events (ranging from 2 to 10 per patient) due to conventional treatment with anti epileptics. By adding CBD, the number of patients reporting adverse events dropped to 7,9% and the number of adverse events reported dropped to an average of 1 (ranging from 0 to 2 per patient). The results are very expressive and advocate in favor of the safety of CBD treatment as an additional therapy (add-on). When we analyze separately each adverse event reported, we have that the CBD reduces the occurrence of fatigue, sleepiness, irritability, insomnia, appetite loss, aggressiveness, nausea, dizziness, anxiety, confusion, weight loss, vomiting and obsessive behavior from 5 to 10 times. The only adverse events that increased with the adding of CBD were weight gain and increased appetite (with an increase of about 2 times). It is worth mentioning that these “adverse events” are not relevant when considering the therapeutic benefits and may even be considered beneficial for some patients, as is the case of patients with anorexia or cachexia provoked by HIV or chemotherapy.

Table XX - Adverse events reported in studies of CBD and conventional anti epileptics. Reproduced from Hussain et al, 2015.

| **Adverse event** | **Before CBD** | **With CBD** |
| --- | --- | --- |
| **Number of AEs** | **5 (2-10)** | **1 (0-2)** |
| **Average** | **36.9%** | **7.9%** |
| Fatigue | 70.1% | 9.4% |
| Sleepiness | 70.9% | 12.8% |
| Irritability | 59.0% | 9.4% |
| Insomnia | 43.6% | 5.1% |
| Apetite reduction | 40.2% | 6.0% |
| Agressiveness | 40.2% | 4.3% |
| Nausea | 35.9% | 6.8% |
| Dizziness | 35.9% | 0.0% |
| Anxiety | 35.9% | 3.4% |
| Mental confusion | 35.0% | 0.9% |
| Weight loss | 35.0% | 4.3% |
| Emesis | 23.9% | 2.6% |
| Obsessive complex | 23.9% | 2.6% |
| Weight gain | 17.9% | 29.1% |
| Rash | 10.3% | 0.0% |
| Apetite gain | 12.8% | 29.9% |

## 5 Conclusion

In conclusion, this meta analysis suggests that treatments using CBD demonstrate efficacy, at least in the population of patients with reported refractory epilepsy and have an adequate safety profile, considering risks and benefits inherent to the treatment of this severe neurological condition. A considerable share of patients obtain benefits from this treatment, and the adverse events, when they occur, are fairly mild.

Apparently, CBD enriched extracts are more potent and have a better profile of adverse effects (but not a greater efficacy) than purified CBD products, at least for this population of patients with refractory epilepsy. The lack of standardization among Cannabis extracts does not allow us to infer directly which characteristics of the product provide this therapeutic advantage, but it probably is related to other compounds present at the formulation which act sinergistically to CBD. Among the known compounds of the plant, we have at least 7 anti convulsives, including CBD. Such compounds are Canabidiol (CBD), Δ9-tetrahydrocanabinol (THC), and other minor canabinoids such as tetrahydrocanabivarin (THCV), canabidivarin (CBDV), canabinol (CBN), and other non canabinoid compounds (terpenes), like linalool and myrcene. These last two are naturally occurring in essential oils of other plants known and used by men like bay leaves and hops, in the case of myrcene, and lavender and citrus, in the case of linalool. Controlled studies with standardized Cannabis extracts are necessary to confirm if these compounds contribute isolated or sinergistically for the anti convulsive effect of Cannabis and its byproducts.

